# Predatory fish preferentially target virtual prey with Lévy motion rather than Brownian motion

**DOI:** 10.1101/2022.07.17.500339

**Authors:** Christos C. Ioannou, Luis Arrochela Braga Carvalho, Chessy Budleigh, Graeme D. Ruxton

## Abstract

Of widespread interest in animal behaviour and ecology is how animals search their environment for resources, and whether these search strategies are optimal. However, movement also affects predation risk through effects on encounter rates, the conspicuousness of prey, and the success of attacks. Here we use predatory fish attacking a simulation of virtual prey to test whether predation risk is associated with movement behaviour. Despite often being demonstrated to be a more efficient strategy for finding resources, we find that prey displaying Lévy motion are twice as likely to be targeted by predators than prey utilising Brownian motion. This result emphasises that costs of predation risk need to be considered alongside the foraging benefits when comparing different movement strategies.

## Introduction

Movement to find resources is a general hallmark of the animal kingdom, displayed by most species in at least some life history stages. Describing the trajectories of these movements has long been of interest to biologists [1]. A very common theoretical description of such trajectories is a random walk, where the trajectory is broken down into a series of steps, and characteristics of steps (specially direction and length) are drawn stochastically from defined probability distributions. Two commonly studied random walks are differentiated by the distribution of step lengths: in Brownian motion this is a negative exponential, while in Lévy motion this is a power-law function that is more heavy-tailed than the negative exponential. In both models most steps are short, but long steps are more common for Lévy than Brownian motion [2,3].

Over the last 25 years Lévy motion has been subject to intense study. Theoretical work suggests that Lévy motion is a highly efficient way to search the environment for hidden resources [4–6], and trajectories from a broad range of species have been suggested to fit Lévy motion [7,8, although see 9]. However, except for apex predators, how animals move in space also determines encounter rates with, and conspicuousness to, their own predators [10–12]. It is also generally considered that unpredictable movement by prey may hinder capture by predators [13–15, although see 16]; the lack of turns during longer step lengths may increase the short-term predictability of movement and so increase predation risk. Here we use a system of fish predators (three-spined sticklebacks, *Gasterosteus aculeatus*) targeting computer-generated prey whose motion can be entirely controlled [17,18] to test the hypothesis that prey with Lévy motion are targeted preferentially relative to prey with Brownian motion, potentially revealing a cost of Lévy motion that counteracts the benefits for finding resources [4–7].

## Methods

### Experimental setup

The experiment used a similar setup to other virtual prey experiments [17,18] but had a larger arena in which the virtual prey could move to increase the spacing between the prey, minimising the prey being perceived as being within the same group. The fish were presented with two-dimensional prey projected onto the longest wall of a glass tank (Figure S1). The trajectory of each prey was simulated using the TrajGenerate function in the Trajr package (version 1.4.0 [19]) in R version 4.1.2 (Figure 1(a)) to create trajectories for prey with a random walk and a distribution of step lengths that followed either Brownian or Lévy distributions (Figure 1(b,c), Figure S2). Each simulation included 5 prey of each movement type, and prey were otherwise identical in appearance. When projected on the screen, the prey appeared as white dots moving at a constant speed (17 mms^-1^) and size (3 mm diameter), and could move within an area of 66 × 28 cm (width × height). Trajectories were long enough to generate 20-minute videos at 60 frames per second. Each trial used a unique simulation of the prey to avoid pseudoreplication.

**Figure 1:**
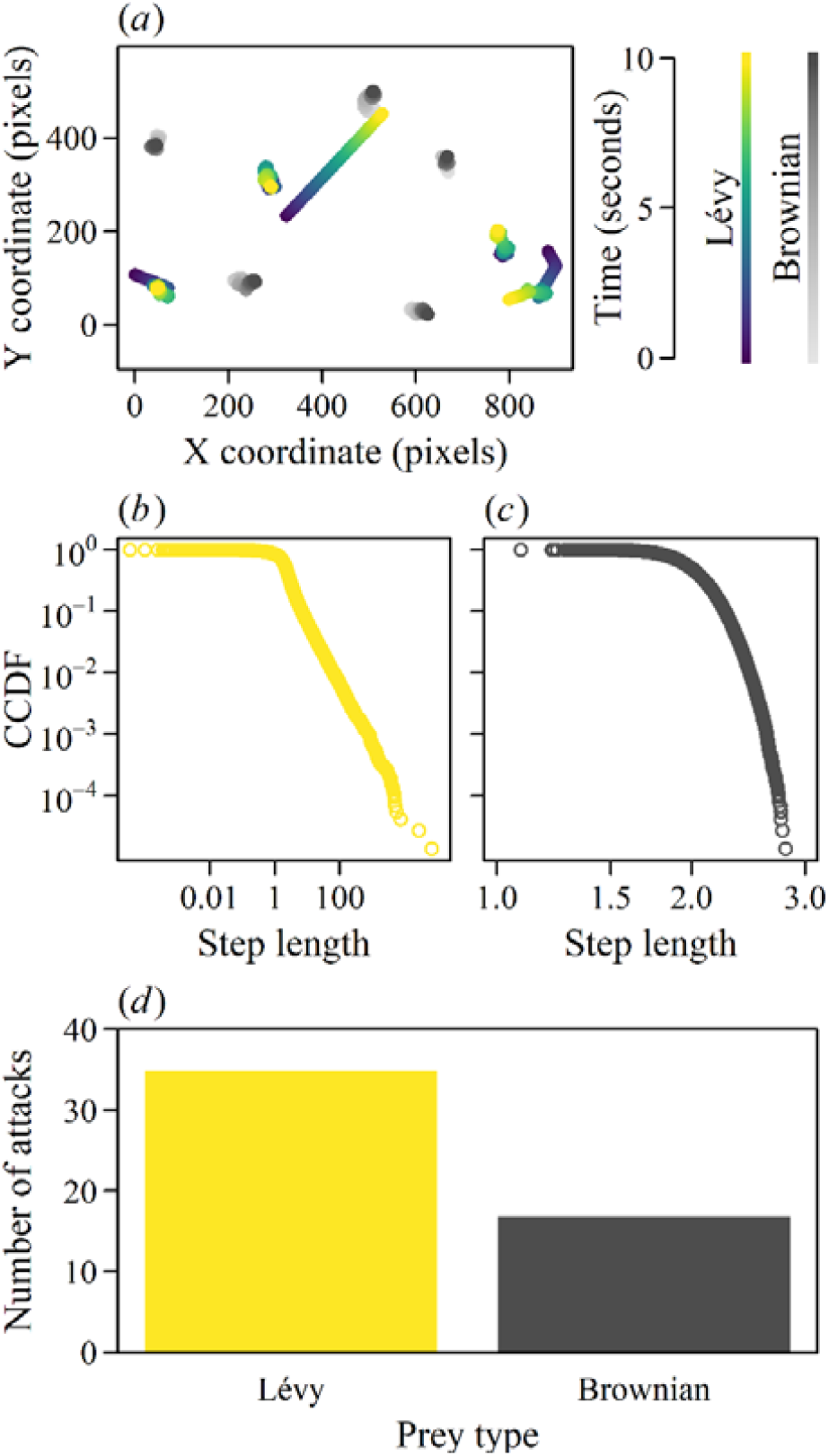
The prey simulation and the number of attacks on each prey type. In (a), the trajectories of the prey over 10 seconds are shown. (b) and (c) show the complementary cumulative distribution function (CCDF) for example Lévy and Brownian trajectories as generated by the Trajr package before these were rediscretised to give a constant speed of movement for each prey (Figure S2). (d) shows the number of trials where the first attack targeted each prey type.

### Experimental procedure

All procedures were approved by the University of Bristol ethical review group (UIN/21/003). 15 fish were haphazardly netted from stock tanks the day before testing and moved to the holding area of the test tank (Figure S1). The next day, the simulation projection and camcorder recording for that trial were started, and a single fish was netted from the holding area to the test arena. Trials were ended when the fish made their first attack or 15 minutes had elapsed without an attack. Each fish was tested only once. Trials were conducted between the 9th and 26th November 2021.

### Data extraction and statistical analysis

From the camcorder recordings, the frame at which the fish made their first attack was identified. Only the first attack was included in the analysis. The identity of the prey (Brownian or Lévy) was determined by visual comparison of which prey was attacked in the video to the plotted prey coordinates from the simulation at the corresponding time step. An exact binomial test was used to test whether the frequency of attacking each prey type differed from the random expectation of 0.5, as the two prey types were equally common.

The difference in Brownian versus Lévy motion is the frequency and duration with which individuals move in a straight line (Figure 1(b,c), Figure S3). We thus used custom randomisation tests to determine whether the turning angle of the targeted prey was greater or less than expected from the predators attacking that prey at randomly chosen times in that trial before the attack (i.e. whether the predators preferentially attacked that prey when they were moving in straighter or more sinuous paths than usual; see figures S3, S4 and S5 for details). A second randomisation procedure tested whether the turning angle of the attacked prey was greater or less than expected if the predators randomly chose a prey to attack at the same time step that the observed attack occurred (figures S3 and S6). The simulations and statistical analysis can be recreated from the R code and experimental data provided as electronic supplementary material.

## Results

In 54 of the 117 trials the test fish made an attack within the 15-minute trial time. Data from two of these trials were lost due to technical malfunctions. In 35 of the remaining 52 trials with attacks (67%), the fish attacked a prey with Lévy motion (Figure 1(d); exact binomial test: P = 0.018).

There was no evidence from the randomisation tests that the fish preferentially targeted the attacked prey when it was moving in a more or less straight path than expected from randomly-timed attacks on that prey (Table S1 and S2, Figure S5). In contrast, there was a statistically significant tendency for the attacked prey to have straighter movement in the time period immediately before the attack than expected if the predators selected a prey randomly at the same moment as the attack (Table 1, Figure S6).

**Table 1:** The quantiles of the observed proportion of turns (mean value across the 52 trials) and observed mean angle turned (mean value across the 52 trials) in the corresponding distributions of expected values if the fish attacked a randomly-selected prey at the same moment as the observed attacks. As two-tailed tests, the quantiles are statistically significant (marked in bold) if they are less than 0.025 (the attacked prey turned less than expected) or greater than 0.975 (they turned more than expected). See also Figure S6.

| Time window (frames) | Quantile of mean observed proportion of turns in expected distribution | Quantile of mean observed mean turn angle in expected distribution |
| --- | --- | --- |
| 60 | **0.014** | **0.018** |
| 120 | **0.024** | **0.023** |
| 180 | 0.027 | **0.013** |
| 240 | **0.019** | 0.028 |
| 300 | **0.012** | **0.018** |

## Discussion

The fish we used as predators preferentially attacked those with Lévy motion twice as often as those with Brownian motion. Our randomisation tests found no evidence that attacks on prey were more likely when they were engaged in straight movements (i.e. with low turning), suggesting that the timing of attacks was not influenced by prey behaviour. However, when the fish did attack, they preferentially targeted those prey that were turning less than other prey; as prey with Lévy motion were more likely to be turning less, this can explain the preferential targeting of prey with Lévy motion. Exploring how general the effects reported here are across predators and contexts would be welcome. The advantage of a virtual prey system allowing for precise control over prey traits and hence minimising confounding effects is, however, countered by the limited range of predators that are suitable for testing in such an artificial, laboratory-based set up. Another valuable extension of the results here would be to explore the consequence of allowing predators repeated exposure to prey that vary in their movement pattern, as recently we have shown that predators can adapt to unpredictability of fleeing prey [16]. Any such future work would ideally not only consider the choice of which prey is targeted amongst those displaying different types of motion, but also allow the prey to be captured and consumed so that attack success can be measured.

Although we find there may be cost to Lévy motion, Lévy motion might still be selected for if it is a sufficiently better search strategy to find resources [4–7]. For cases where there is a maximum rate of resource use (for example limited by rates of digesting food [20]), a reduced time searching for resources using efficient, but potentially higher risk, Lévy search would allow for more time spent safe in refuges. Future work could explore the conditions that affect the trade-off between resource acquisition and predation risk, including how the total predation risk and how it changes over time affects movement trajectories.

Another extension of this study would be to explore how movement behaviour of prey interacts with other aspects of their anti-predatory defence, such as living in groups [21]. Camouflage is a very common anti-predator trait [22], and movement is known to adversely affect camouflage [12]. However, it remains unexplored how details of movement (e.g. speed, predictability, rate of change in direction) quantitively affect ease of detection by predators, and how consistent such trends would be across different types of camouflage. However, as well as camouflage, there is mounting evidence that the appearance of moving prey can influence the ability of predators to estimate the speed and direction of prey [23,24]. It may be that patterning in prey might modify the targeting preferences explored here and/or any effects on movement pattern on the subsequent success of attacks by predators.

Previous consideration of Lévy motion has focused almost exclusively on the consequences for foraging. Here we show that there may be consequences for predation risk also. Since most foragers have to contend with potential predators, we hope that our work encourages a broader perspective in future study of Lévy motion patterns in particular, and search behaviour in animals more generally.

## Supporting information

Experimental data

R script for simulations and analysis

## Acknowledgements

This work was funded by a NERC Independent Research Fellowship (NE/K009370/1) and a Leverhulme Trust grant (RPG-2017-041 V) awarded to C.C.I.

