## Supplementary tables and figures for "Predatory fish preferentially target virtual prey with Lévy motion rather than Brownian motion"

1	Table S1, S2

2	Figure S1, S2, S3, S4, S5, S6

3	Supplementary Dataset

4	Supplementary R code

5	References

Table S1: The number of trials where the turning behaviour of the prey immediately preceding when it is attacked is less or more than expected if the predator attacked the same prey at a randomly determined time in the simulation. The turning behaviour of each prey is measured as both the proportion of frames where a turn occurs (see Figure S3) or the mean turning angle. See figure S4 for detail regarding the randomisation. As the number of the 52 trials outside of the expected 95% confidence intervals are low, there is little evidence that the attacks by the fish were timed to target prey when they were moving in straighter or more sinuous paths than expected from randomly-timed attacks (also see figure S5).

| Time window (frames) | N trials observed proportion of turns <0.025 of expected distribution | N trials observed proportion of turns >0.975 of expected distribution | N trials observed mean turn angle <0.025 of expected distribution | N trials observed mean turn angle >0.975 of expected distribution |
| --- | --- | --- | --- | --- |
| 60 | 0 | 4 | 0 | 2 |
| 120 | 0 | 0 | 0 | 0 |
| 180 | 0 | 1 | 0 | 0 |
| 240 | 0 | 1 | 0 | 1 |
| 300 | 1 | 2 | 0 | 2 |

Table S2: The quantiles of the observed proportion of turns (mean value across the 52 trials) and observed mean angle turned (mean value across the 52 trials) in the corresponding distributions of expected values if the fish predators attacked the same prey but at a randomly determined time within each trial. See figure S4 for detail regarding the randomisation. There is no evidence that the attacks by the fish were timed to target prey when they were moving in straighter or more sinuous paths than expected from randomly-timed attacks.

| Time window (frames) | Quantile of mean observed proportion of turns in expected distribution | Quantile of mean observed mean turn angle in expected distribution |
| --- | --- | --- |
| 60 | 0.254 | 0.228 |
| 120 | 0.375 | 0.253 |
| 180 | 0.338 | 0.202 |
| 240 | 0.347 | 0.288 |
| 300 | 0.295 | 0.274 |

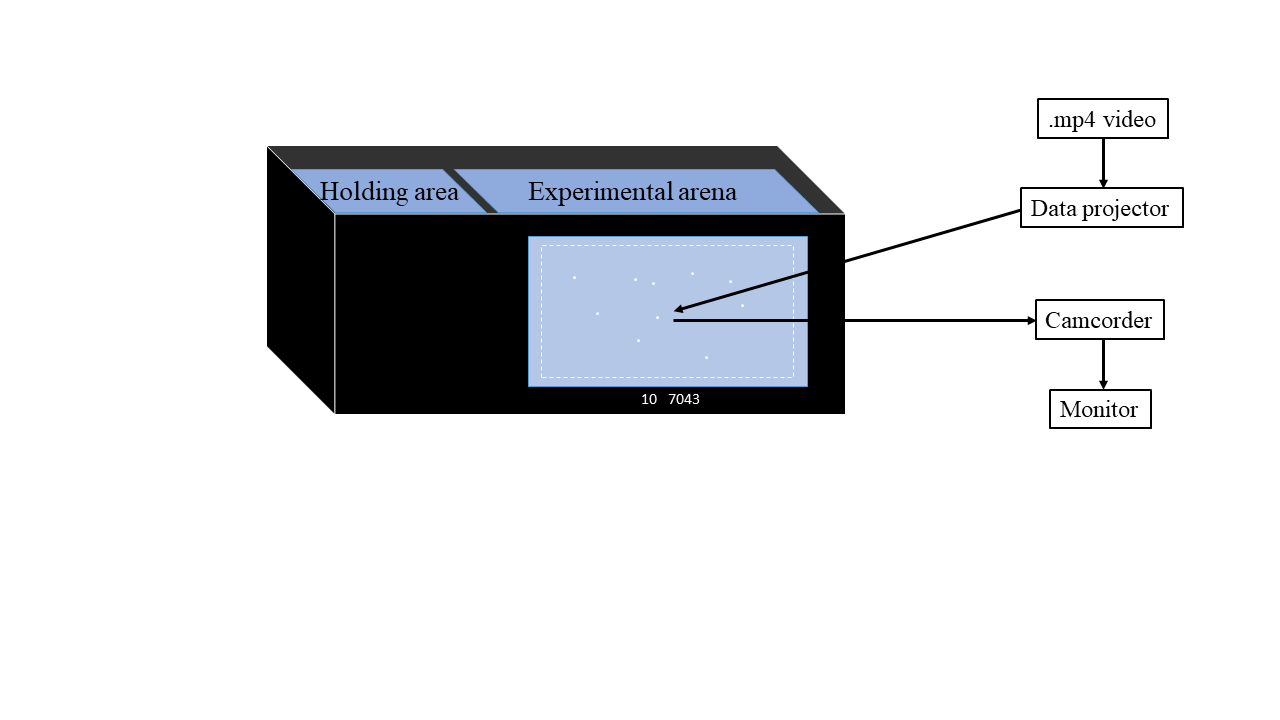

Figure S1: Schematic of the experimental set up, not to scale. Three-spined sticklebacks were obtained from Carp Co. (www.carpco.co.uk) in September 2021. They were kept in 40 × 70 × 34 cm (width × length × height) glass tanks with a flow-through recirculation system, plastic tubes as shelter and plants for environmental enrichment. They were kept at 14°C under a 11:13 light : dark cycle and each tank housed a maximum of 100 fish. All subjects in the holding tanks were fed small granular pellets of fish feed every morning except for days of testing when they were fed after the trials were finished. The 36 × 121 × 47 cm (width × length × height) experimental tank was filled with aged tap water which was continually filtered with an external Eheim Classic 350 filter and chilled with an aquarium chiller (D-D DC300) to maintain the temperature between 13 and 14ºC. The test tank was divided into a 36 × 28 × 47 cm holding area with artificial plants where the fish were habituated the evening before testing, and a 36 × 93 × 47 cm experimental arena where the trials were carried out. A sheet of white translucent plastic film (Rosco gel no. 252) was taped onto the inside of the longest side of the tank facing the projector and camcorder. Black plastic on the outside of the tank framed the translucent screen to create a 84 × 34 cm (width × height) area for the prey to be projected on to. The white dashed box in the figure, which was not projected during the trials, indicates the area within which the prey could move. The video number and time step of the simulation were included under the area in which the prey could move to be visible in the view of the camcorder but out of view of the fish. The other walls of the tank were covered internally with black opaque plastic. A strip light was installed above and behind the back wall of the tank to provide illumination so that the fish was visible through the translucent screen, facilitating the detection of attacks by the fish. A data projector (BenQ MW523) was positioned 132.5 cm in front and above the front tank wall and a camcorder was positioned 104.5 cm directly in front of the front wall and below the projector, filming at 1920 × 1080 and 50 frames per second. Due to technical difficulties, Panasonic VX870 and then Panasonic HC-X920 camcorders were used to record the trials. The camcorder was connected to a monitor to remotely observe the trials. Black sheets enclosed the space between the tank and the projector and camcorder in order to provide a dark background against which the fish could view the prey, and to minimise external disturbances.

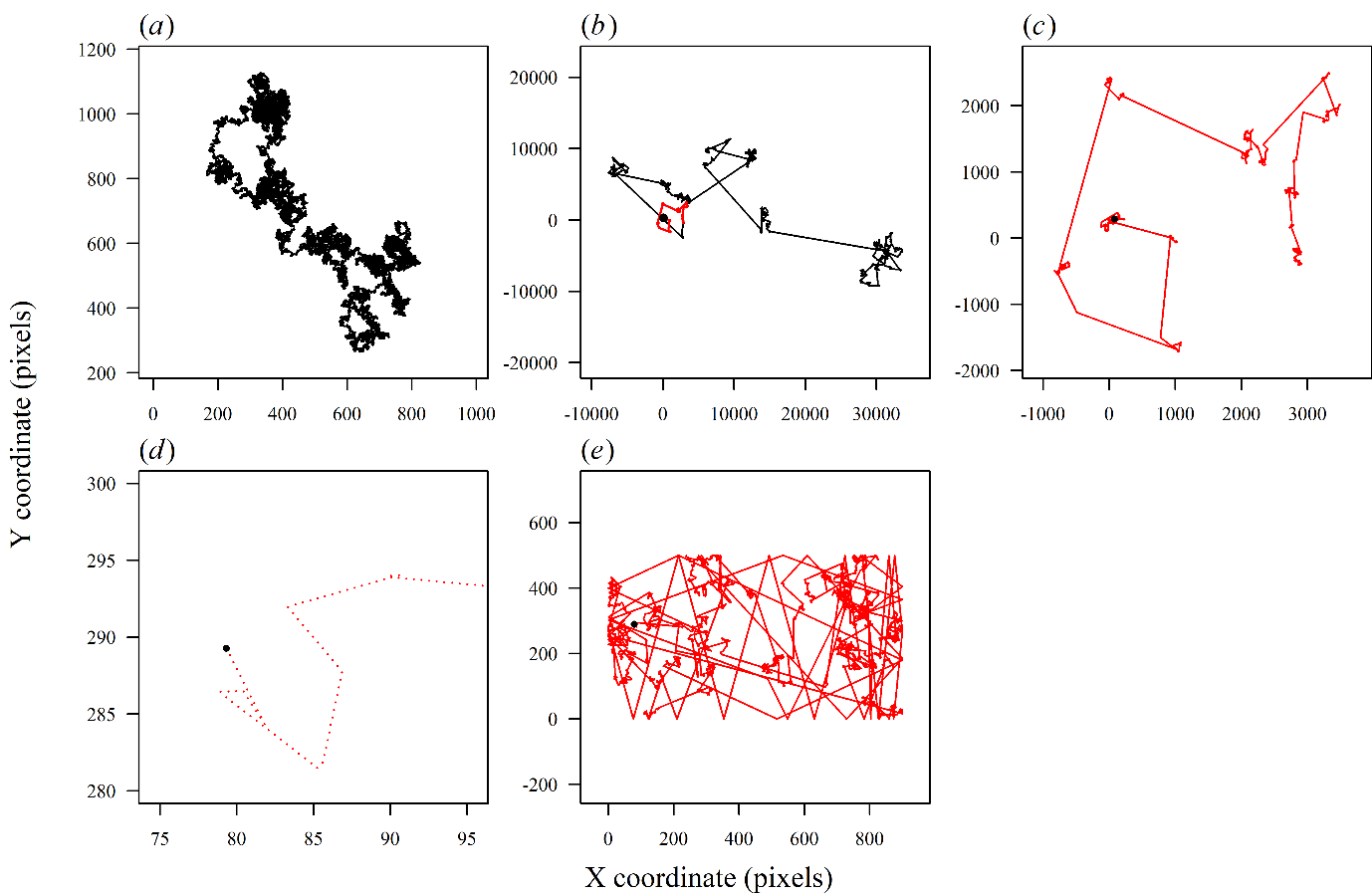

Figure S2: Creating the simulated prey trajectories. The prey are simulated in two dimensions to move with Brownian or Lévy motion using the TrajGenerate function from the R package Trajr, version 1.4.0 [1]. Example Brownian and Lévy trajectories are displayed in panels (a) and (b), respectively; note the greater range of x and y coordinates in the Lévy example. To avoid pseudoreplication, a unique random seed was set before each path was generated, ensuring that no two paths were identical. The starting position of the prey at the beginning of each video (indicated by the filled black circle) was randomly drawn from a uniform distribution of x coordinates between 50 and 850 pixels, and a uniform distribution of y coordinates between 50 and 450 pixels. Again, a unique random seed was set before each random draw, ensuring the starting positions were unique for each prey and that the trajectories could be reconstructed exactly. The TrajRediscretize function in Trajr version 1.4.0 was then used to resample points along each trajectory with an equal distance between each pair of points, causing the prey to move at a constant speed. Panel (c) demonstrates the rediscretised Lévy trajectory from (b); the segment of trajectory in (b) that corresponds to (c) is coloured red. The rediscretised trajectory has the same number of steps as the original trajectories, but as the rediscretised step lengths (d) are shorter than the original step lengths, the rediscretised trajectories are shorter. Reflective boundary conditions were then imposed to ensure the prey were restricted in their x coordinate between 0 and 900 pixels and y position between 0 and 500 pixels. (e) shows the trajectory from (c) after the reflective boundary conditions were applied. At each time step in the simulation, each prey’s x and y coordinates were plotted as scatter plots. The saveVideo function from the animation package version 2.6 [2] was used to convert these scatterplots into a .mp4 format video at a resolution of 1280 × 800 and 60 frames per second, matching the resolution of the data projector.

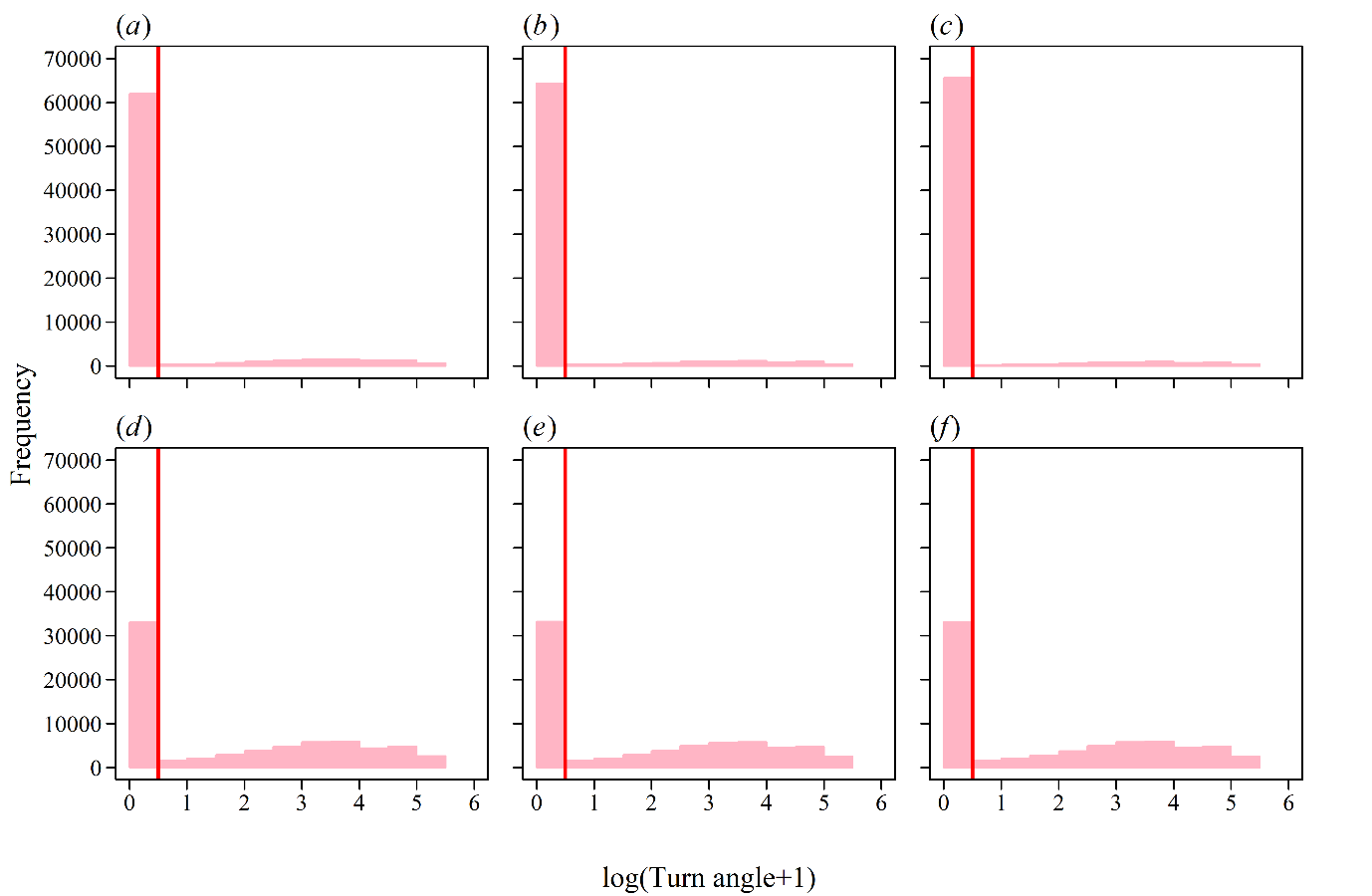

Figure S3: Example distributions of turning angles of single prey over the duration of the simulations. (a, b, c) show turning angles from the prey following Lévy motion that were attacked in trials 2, 4 and 6 (respectively), and (d, e, f) show turning angles from the prey following Brownian motion that were attacked in trials 1, 3 and 5 (respectively). As expected, prey with Brownian motion have greater turning angles than prey with Lévy motion. The turning angles are log(turning angle+1) transformed to illustrate the bimodal distribution of the data, where a threshold can be applied at 0.65 degrees (0.5 in the log(turning angle+1) axis, indicated by the vertical red line) to differentiate turns from movements that can be considered movement in a straight line.

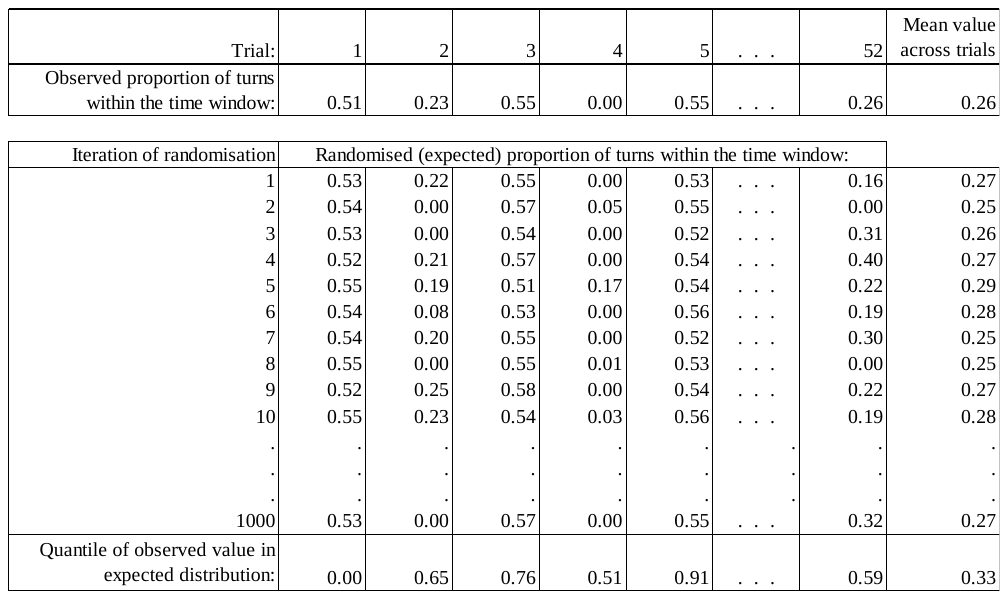

Figure S4: Schematic of the two randomisation approaches to test whether the observed attacks on prey occurred at times when the target prey was turning more or less than usual. In the first test, for each trial (each column) the observed turning value is compared with the 1,000 iterations of the randomisation where the turning angle is calculated for the same sized time window but beginning at a randomly determined time (in the period from the start of the trial until the observed attack is made). This yields the quantile of the observed turning value relative to the expected distribution for each trial (final row). The second approach uses the mean values for the turning variable across the 52 trials (final column). Here, the observed mean turning value is compared to the distribution from 1,000 mean values expected if the attacks were made at the same randomly selected times as in the first approach. The two turning variables are the proportion of turns within the time window that are greater than 0.65 degrees (Figure S3; values in the figure above show an example data set of this variable) and the mean turning angle within that time window. The procedure is repeated for five time windows across which the turning variables are calculated (60, 120, 180, 240 and 300 frames, equal to 1, 2, 3, 4 and 5 seconds).

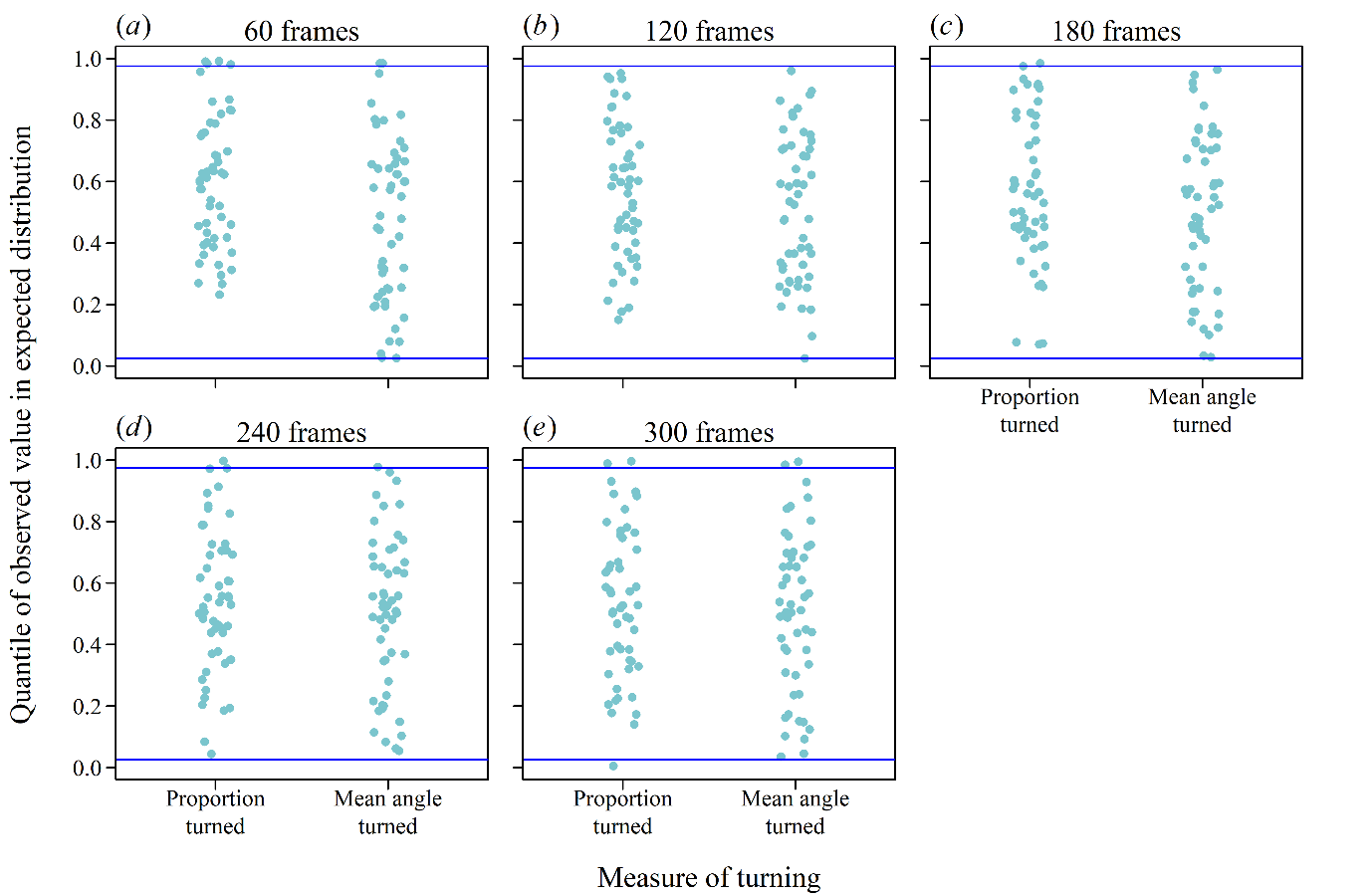

Figure S5: Results of randomisation testing whether the timing of attacks on prey is dependent on the attacked prey’s turning behaviour in the time window immediately preceding an attack. The turning behaviour of each prey is measured as both the proportion of frames where a turn occurs (see Figure S3) or the mean turning angle. This analysis is conducted on each trial separately: the observed turning value is compared to the same statistic from 1,000 randomly selected time points from the start of the trial until the moment of attack. The quantile of the observed value within the corresponding expected values is shown as blue dots (with jitter added to aid visualisation). Quantiles smaller than the lower 2.5% interval (0.025, indicated by the lower solid horizontal line) indicate the observed attack occurred on prey turning less than expected from a randomly-timed attack, and quantiles greater than the upper 97.5% (0.975, the upper solid horizontal line) indicate the attacked prey turned more than expected. The procedure is repeated for five time windows across which the turning variables are calculated (60 (a), 120 (b), 180 (c), 240 (d) and 300 (e) frames, equal to 1, 2, 3, 4 and 5 seconds).

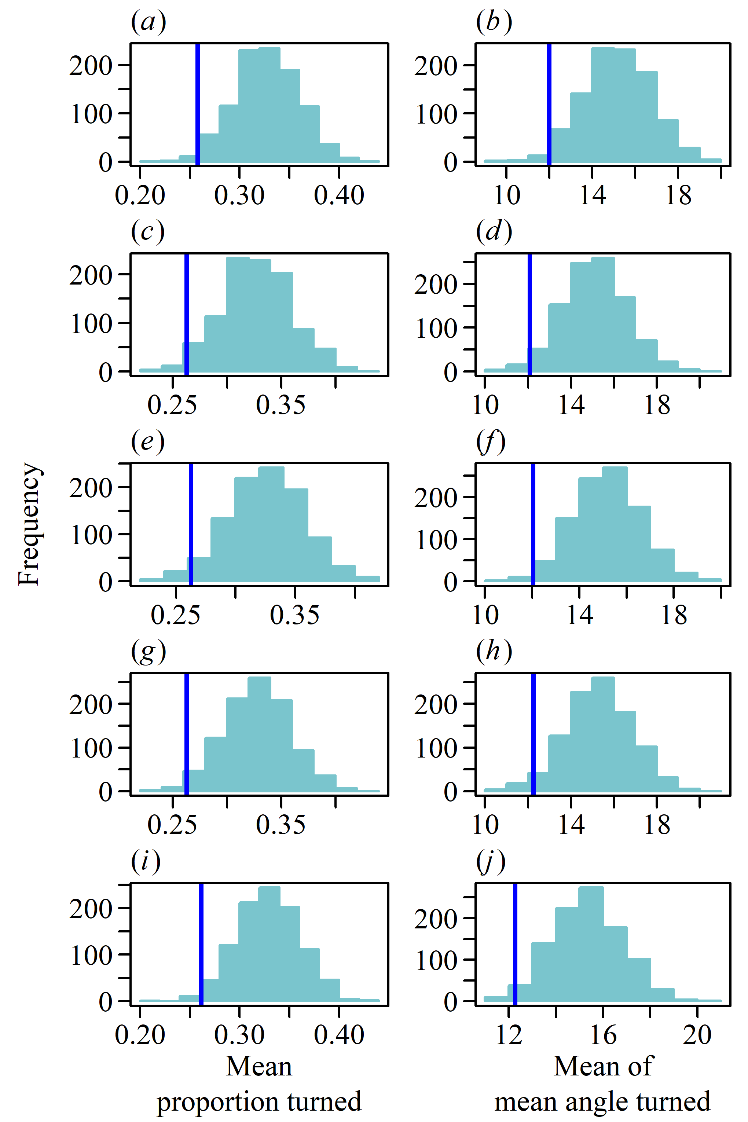

Figure S6: Results of randomisation testing whether the choice of which prey to attack is dependent on the prey’s turning behaviour in the time window immediately preceding the observed attacks. The turning behaviour of each prey is measured as both the proportion of frames where a turn occurs (left column) or the mean turning angle (right column). The dark blue vertical line shows the mean turning value (averaged (mean) across trials) for the observed attacks in the experiment. The light blue histograms show the expected distribution of these averaged turning values if the predator chose to attack prey (at the same moment as the observed attacks) at random, i.e. with equal probability regardless of their turning behaviour. 1,000 iterations of the randomisation are conducted at each time window to generate the expected distributions of turning values. Quantiles for the observed mean values smaller than the lower 2.5% interval indicate the attack prey was turning less than expected if the predator chose each prey to attack with an equal probability. The procedure is repeated for five time windows across which the turning variables are calculated (60 (a, b), 120 (c, d), 180 (e, f), 240 (g, h) and 300 (i, j) frames, equal to 1, 2, 3, 4 and 5 seconds).

Supplementary Dataset: The data from each trial resulting in an attack. This includes the when in the video the attack occurred and which prey was attacked (prey 1 to 5 have Brownian motion, prey 6 to 10 have Lévy motion).

Supplementary R code: The R code used to create the simulations and analyse the data from the experimental trials.
